# Complete mitochondrial genome sequence of Bos frontalis (Gayal) from Bangladesh

**DOI:** 10.1101/2020.12.31.424938

**Authors:** GK Deb, R Khatun, SMJ Hossain, SS Rahman, MAB Bhuiyan, S Mobassirin, S Afrin, M Billah, A Baten, NR Sarker, MSA Bhuyian, AMAMZ Siddiki

## Abstract

The Gayal is a large-sized endangered semi-domesticated bovine species belonging to the family Bovidae, tribe Bovini, group Bovina, genus Bos, and species Bos frontalis. It is also called the Mithan or Mithun. Mitochondrial genome is considered as an important tool for species identification and monitoring the populations of conservation concern and therefore it becomes an obligation to sequence the mitochondrial genome of Bagladeshi gayal. We want to identify some important genes related to a particular trait such as those associated with adaptation, muscle strength, or prolificacy. The data will help explore evolutionary relationships with closely related species. The mitogenome of *Bos frontalis* is 16,347 bp in length and nucleotide composition is AT-based (60.21%), contains 37 genes including 13 protein-coding genes, 22 tRNA genes, 2 rRNA genes, and a control region.

## Introduction

Gayal could be a semi-wild animal and originated from the cross between Gaur and domestic cattle. The gayal originated within the hilly areas of Bhutan, eastern India, eastern Bangladesh, northern Myanmar, and northwestern Yunnan, China with elevation up to 3000 m. within the soggy jungle of hills, gayal is a free-ranging animal without arranged mating. The stereotypical length of the body and head is 2.5 to 3.3 meters; the tail extent ranges from 0.7 to 1.05 meters. The shoulder zenith is 1.65 to 2.2 meters. A pair of horns are present in both sexes; horn length ranges from 0.6 to 1.15 meters. The hair of B.frontalis is dark reddish-brown to blackish brown, with white stockings. Adult males are about 25% larger and heavier than females. A characteristic hump of raised muscle may be seen over the shoulders; this can be the results of elongated spinal processes on the vertebrae [1]. it’s sometimes described as a semi-domestic animal. In Bangladesh, it’s found only within the Bandarban Hill district [2]. Gayal (Mithun) is a very profitable grazer on steep hilly slopes differentiated from other animals. Gayal is primarily stern as a meat animal and highly offered among the tribal people of the north-eastern region of India. Nevertheless, gayal is additionally used as a ceremonial animal and plays a crucial role within the economical, gregarious, and folklife of the tribal people of north-eastern India.

Phylogenetic experiments in multiple studies constructed on mtDNA or Y-chromosomal DNA place Gayal in a clash clustering place with relevant cattle, Zebu, and wild Gaur. for example, Chinese Gayal, or Dulong cattle, are familiar to harbor Zebu or Taurine mtDNA footprints, suggesting hybrid origin, and more studies have shown a high mtDNA and Y-chromosomal DNA sequences similarity between Gayal and Guar. One study has even placed the Gayal as a definite and separate species/subspecies. Experimentation has disclosed a high genomic branching among bovine species. In consequence, mapping of resequencing data from 1 bovine species onto the testimonial genome of various species (for example, gayal vs cattle) produces a path for biases and/or errors within the sequence calibration and SNP calling approach [3]. To precise, this whole study came up with the knowledge of Bos frontalis mitogenome, which can even be valuable to try and do further taxonomic classification, phylogenetic reconstruction, and implementing conservation strategies Here we reported the complete mtDNA sequences of Bos frontalis from the Hill district of Chittagong.

## Methodology

The computational framework used for this study is represented in Figure 1. A fresh sample of adult male *Bos frontalis* was collected from Rangunia, Chittagong, Bangladesh (Longitude / Latitude - 22.4701 N 92.0286 E), and transported to the laboratory to preserve at 70° C, for further analysis during 2019, December. The blood tissue was dissected and stored with 95% ethanol. Later, high molecular weight genomic DNAs were isolated and purified using the AddPrep Genomic DNA extraction kit (AddBio, Korea) for future evaluation of the quality and quantity of the DNA. Purified DNA was sent for library preparation and been sequenced through commercial suppliers. DNA was sequenced using Illumina NextSeq 500 platform from BGI, China. All the methods had been performed in accordance with the “Regulations for Animal Experiments in Chittagong Veterinary and Animal Sciences University’s unique feature, the Indian and GOB ethical clearance” as required. The mitochondrial genome reads were separated from the whole genome sequence by mapping it against the reference Gayal mitochondrial datasets (MW092171)using SAMTOOLS. The organelle assembler NOVOPlasty V.2.7.2 [4] was used to assemble the clean reads. Basic Local Alignment Search Tool (BLAST) database searches were performed to make sure that correct target sequences were obtained. For structural and functional annotation of the mitogenome two web based tools- MITOS [5] and GeSeq [6] were used respectively. Transfer RNA genes were identified by tRNAscan-SE [7] which default search mode using vertebrate mitochondrial genetic code for tRNA structure prediction. Another tool, OGDRAW was used to construct the circular representation of the entire mitogenome [8]. Finally, mtDNA sequences were aligned and a phylogenetic tree was constructed by using MEGA X [9].

**Figure 1.**
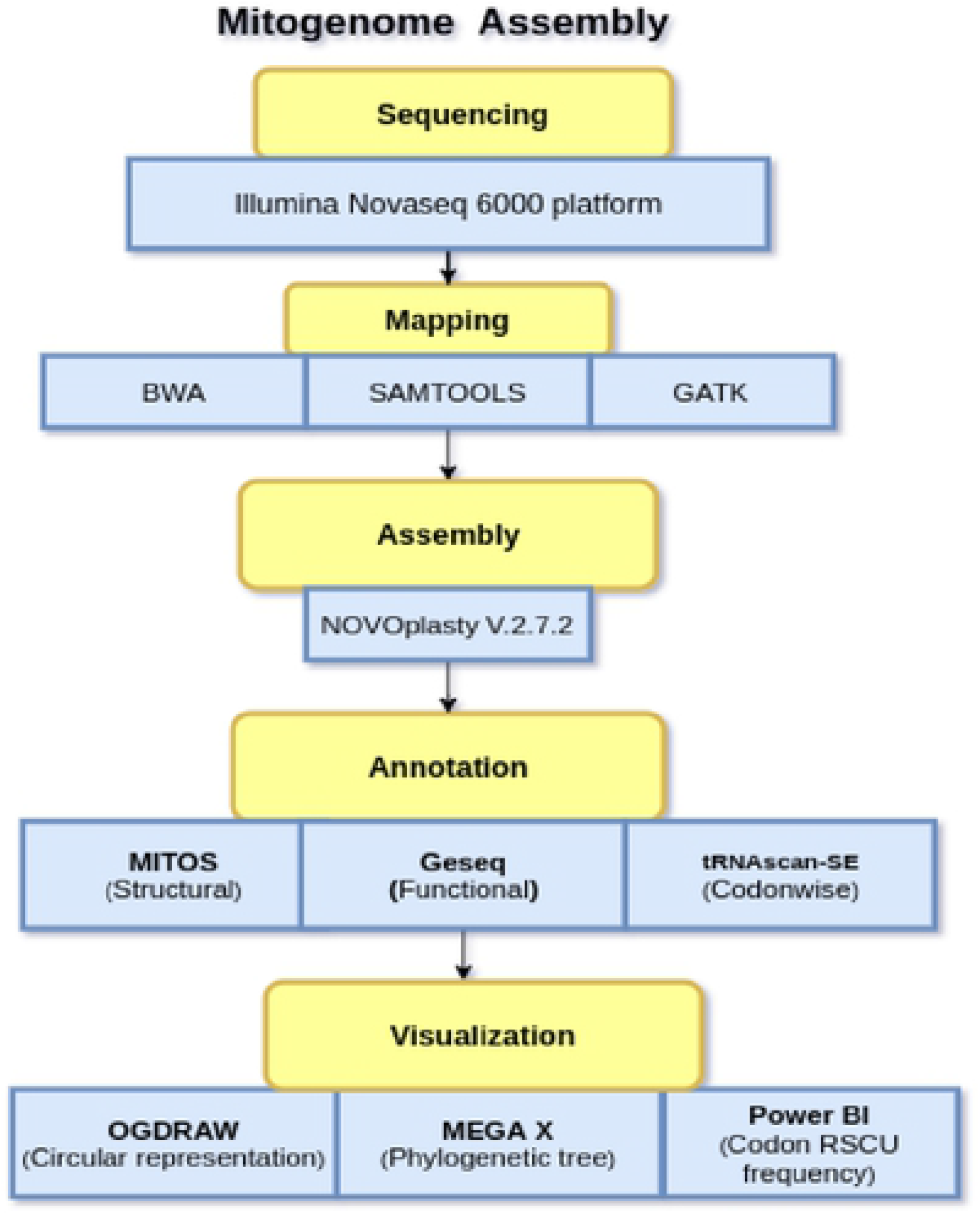
The computational framework used for Bos_frontalis metagenomic analysis where the methodology is divided into 5 steps.

## 3. Result and discussion

### 3. 1. Genome organization and base composition

The complete mitochondrial genome of *B*.*frontalis* is a closed circular and double-stranded molecule of 16,347 bp in size, containing 13 PCGs (cox1-3, nad1-6, nad4L, cob, atp6, and atp8), 22 tRNA genes (one for each amino acid and two each for leucine and serine), 2 rRNA genes (rrnS or 12S and rrnL or 16S) and a major non-coding region known as the control region (CR). Among these 37 genes, 28 genes are encoded by the heavy (H) strand whereas the light (L) stand encodes the remaining nad6 and 8 tRNAs (Table 1; Fig. 2). Therefore, no significant variation is noticed in the length of the PCGs, rRNAs and tRNAs within the mitogenome of different *Bos* species (Fig. 3). This finding advocates the consistent nature of the genome size and gene organization of Bangladeshi *Bos frontalis* with respect to the mitochondrial genomes of Indian Mithun and Gaur that have been reported earlier [10–12]. The AT and GC content of the mitochondrial genome is observed to be 60.21% and 39.79% respectively, which indicates that the nucleotide composition is overall biased towards adenine and thymine (Fig. 4). Similar trend has frequently been observed among the previously reported *Bos* species [11–15].

**Table 1.** Summarized characteristic features of the mitochondrial genome of Bangladeshi *Bos frontalis*. The (+) and (−) values in the intergeneric spacer column represent intergeneric nucleotides and overlapping regions between the genes respectively.

| Genes | Strand | Position | Size (bp) | Intergenic spacer (bp) | Anticodon |
| --- | --- | --- | --- | --- | --- |
| ATP6 | H | 10-684 | 675 | 5 |  |
| COX3 | H | 690-1472 | 783 | 1 |  |
| trnG | H | 1474-1542 | 69 | 0 | TCC |
| ND3 | H | 1543-1887 | 345 | 2 |  |
| trnR | H | 1890-1958 | 69 | 0 | TCG |
| ND4L | H | 1959-2252 | 294 | -3 |  |
| ND4 | H | 2249-3616 | 1368 | 10 |  |
| trnH | H | 3627-3696 | 70 | 0 | GTG |
| trnS1 | H | 3697-3756 | 60 | 1 | GCT |
| trnL1 | H | 3758-3827 | 70 | 6 | TAG |
| ND5 | H | 3834-5630 | 1797 | 7 |  |
| ND6 | L | 5638-6156 | 519 | 3 |  |
| trnE | L | 6160-6228 | 69 | 4 | TTC |
| CYTB | H | 6233-7366 | 1134 | 9 |  |
| trnT | H | 7376-7445 | 70 | 0 | TGT |
| trnP | L | 7445-7510 | 66 | 0 | TGG |
| Control region | H | 7511-8431 | 921 | 0 |  |
| trnF | H | 8432-8498 | 67 | 0 | GAA |
| rrnS | H | 8499-9454 | 956 | 0 |  |
| trnV | H | 9455-9521 | 67 | -1 | TAC |
| rrnL | H | 9520-11089 | 1570 | 1 |  |
| trnL2 | H | 11091-11165 | 75 | 8 | TAA |
| ND1 | H | 11174-12118 | 945 | 5 |  |
| trnI | H | 12124-12192 | 69 | -2 | GAT |
| trnQ | L | 12190-12261 | 72 | 2 | TTG |
| trnM | H | 12264-12332 | 69 | 0 | CAT |
| ND2 | H | 12333-13370 | 1038 | 4 |  |
| trnW | H | 13375-13441 | 67 | 1 | TCA |
| trnA | L | 13443-13511 | 69 | 1 | TGC |
| trnN | L | 13513-13585 | 73 | 32 | GTT |
| trnC | L | 13618-13684 | 67 | 0 | GCA |
| trnY | L | 13685-13752 | 68 | 1 | GTA |
| COX1 | H | 13754-15292 | 1539 | 3 |  |
| trnS2 | L | 15296-15364 | 69 | 7 | TGA |
| trnD | H | 15372-15439 | 68 | 1 | GTC |
| COX2 | H | 15441-16121 | 681 | 6 |  |
| trnK | H | 16128-16194 | 67 | 1 | TTT |
| ATP8 | H | 16196-16345 | 150 | - |  |

**Figure 2.**
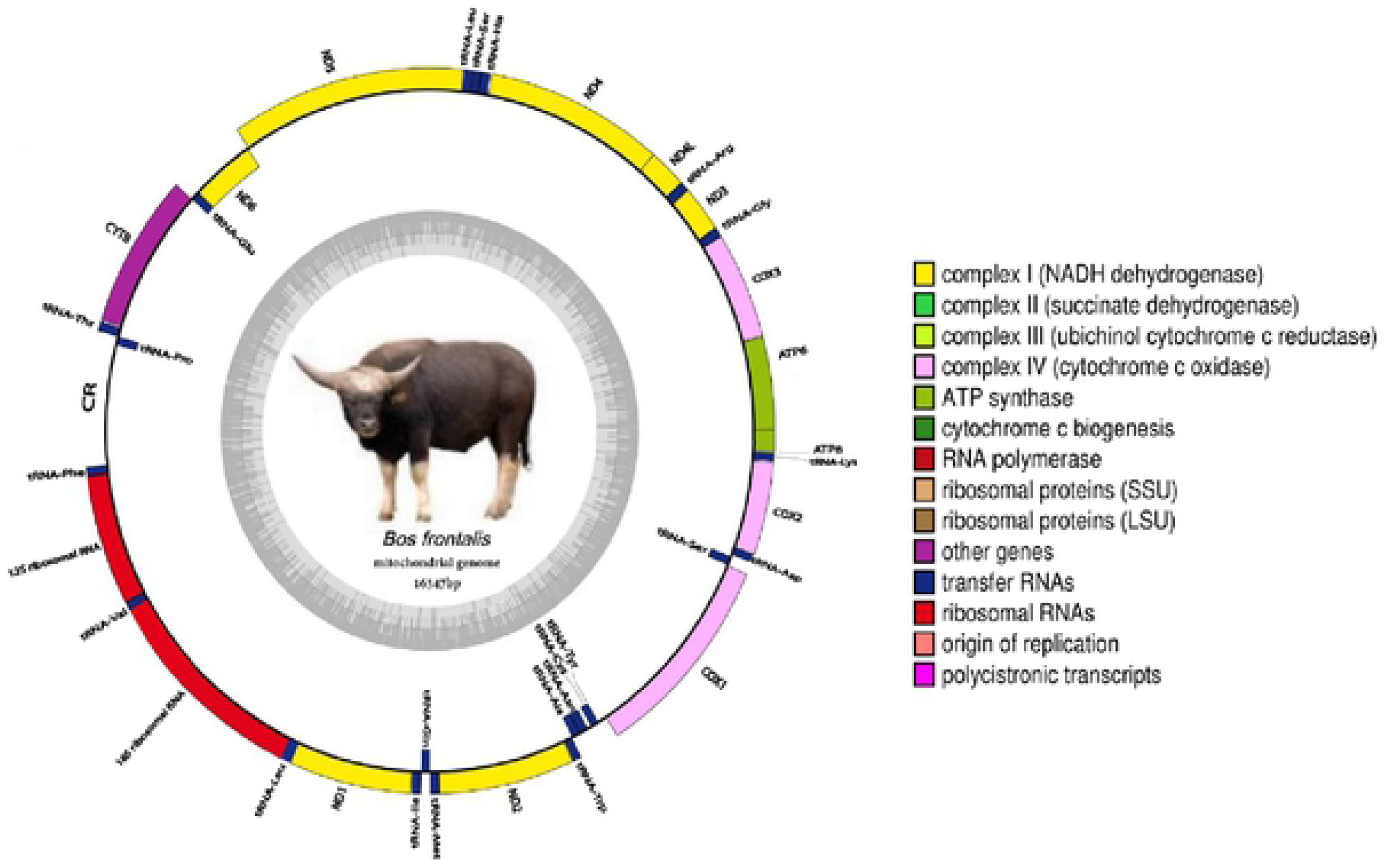
Circular map of the complete mitochondrial genome of the *Bos frontalis*. The colored blocks outside the circle denote 28 genes encoded on the H-strand whereas the colored blocks inside the circle denote the remaining 9 genes encoded on the L- strand. The total GC content of the mitochondrial genome is represented by an inner ring (grey color). The mitogenome map is generated by the webserver OGDRAW [8].

**Figure 3.**
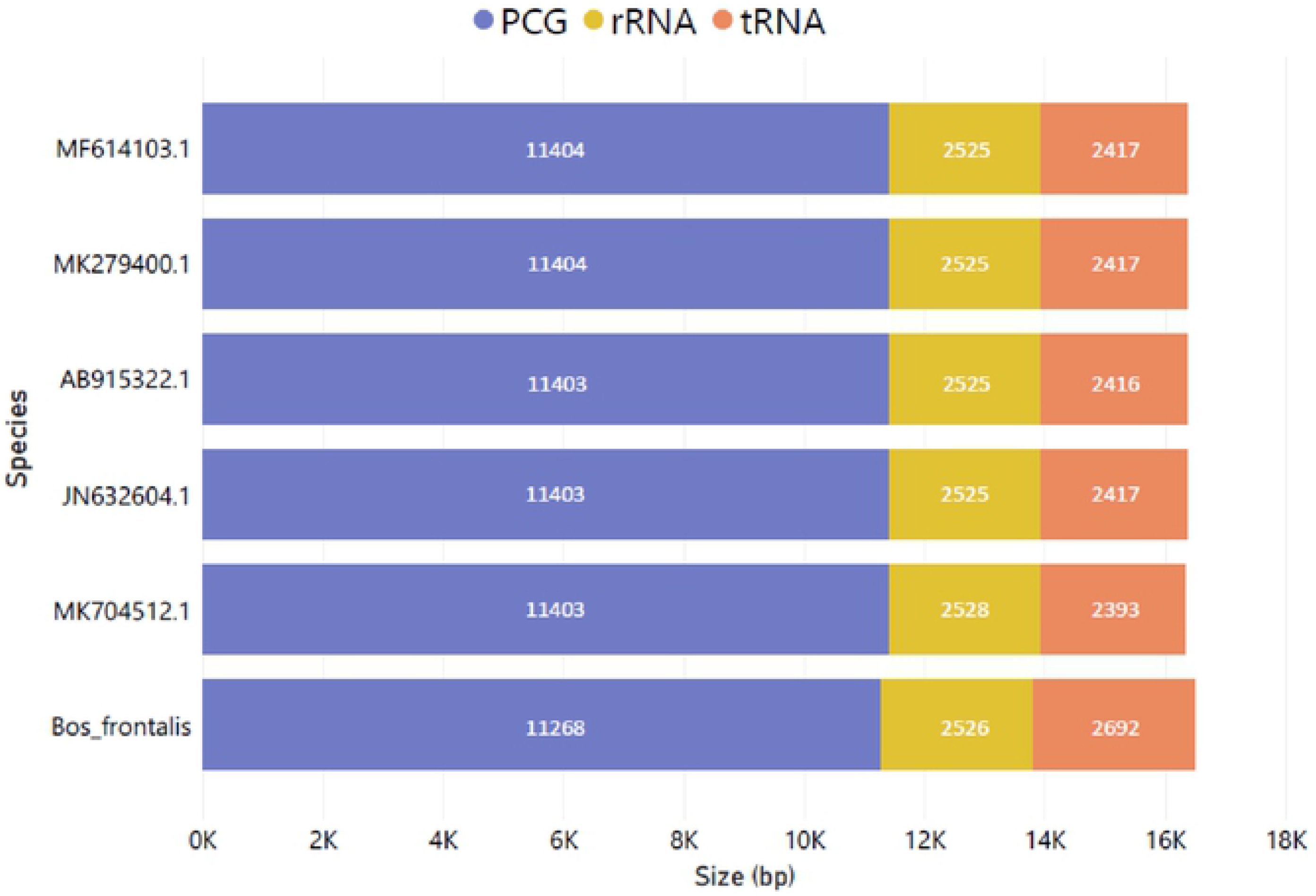
The size of the PCGs, rRNAs, and tRNAs respectively among the closely related *Bos* species.

**Figure 4.**
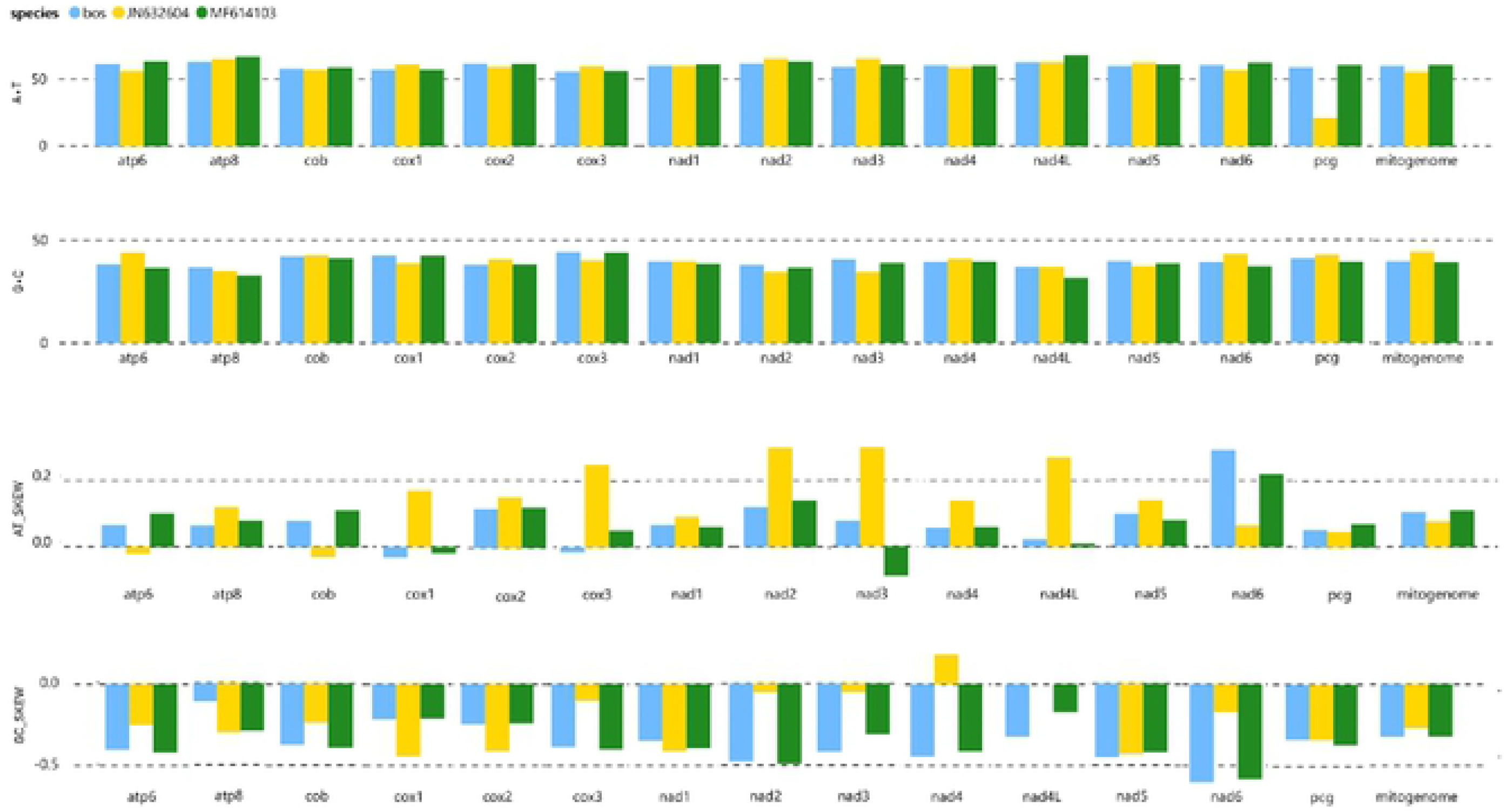
The AT and GC content and skewness of the Bangladeshi *Bos frontalis* and two other *Bos* species.

Moreover, the mitochondrial genome shows positive AT (0.105), and negative GC (−0.322) skews (Fig. 4) which suggests the higher content of adenine and cytosine than their respective complementary nucleotides guanine and thymine. In total there are 5 overlapping nucleotides in the range from 1 to 3 bp, which are found at 3 distinct locations. The largest overlapping region (3 bp) is observed between the two protein-coding genes NADH dehydrogenase subunit 4L (nad4L) and NADH dehydrogenase subunit 4 (nad4). In addition to these, it contains a total of 118 bp of intergenic spacer (IGS) sequence, which is interspersed at 24 regions across the mitochondrial genome with varying ranges from 1 to 32 bp. The longest spacer sequence (32bp) is located between the two tRNA genes trnN and trnC. The putative control region of the mitochondrial genome is located between the tRNA-P and tRNA-F consisting of 921 bp in length. Such similarity has also been observed in other closely related bovine species including Indian Mithun and *B.gaurus* [11–12].

### 3. 2. Protein-coding genes (PCGs) and codon usage

The mitochondrial genome encodes 13 PCGs consisting of 11268 bp in length that accounts for 68.93% of the total mitochondrial genome. The AT and GC content is 58.92% and 41.08% respectively (Fig. 4) indicating the nucleotide compositional biases of the PCGs towards adenine and thymine. Moreover, the positive AT (0.0517) and negative GC (−0.339) skews of PCGs reveals that adenine content is relatively higher than thymine while cytosine content is higher than guanine. Exceptionally cox1 and cox3 genes have slightly negative AT skew while all other PCGs have positive AT skew. Being repeatedly observed in other bovine and mammalian species [11, 15–17], the PCGs are subdivided into 7 NADH dehydrogenase subunits (nad1, nad2, nad3, nad4, nad4L, nad5, and nad6), 3 cytochrome c oxidase (cox1, cox2, and cox3), 2 ATPase subunits (atp8 and atp6) and 1 cytochrome b gene (cob). The size of the PCGs varies remarkably with atp8 (150bp) being the smallest and nad5 (1797bp) being the largest among all. Moreover, a common gene positioning is observed among the four adjacent pairs of PCGs (atp8-atp6, atp6-cox3, nad4L-nad4, and nad5-nad6) likewise the vertebrates [11,16–17].

After excluding the stop codons, the relative synonymous codon usage (RSCU) is calculated and is summarized in Figure 5. The RSCU analysis shows the highest usage of GCU(A), GUA(V), and AAA(K) codons. Furthermore, the most frequently used amino acids in the mitochondrial proteins are proline, threonine, leucine1, asparagine, and serine sequentially.

**Figure 5.**
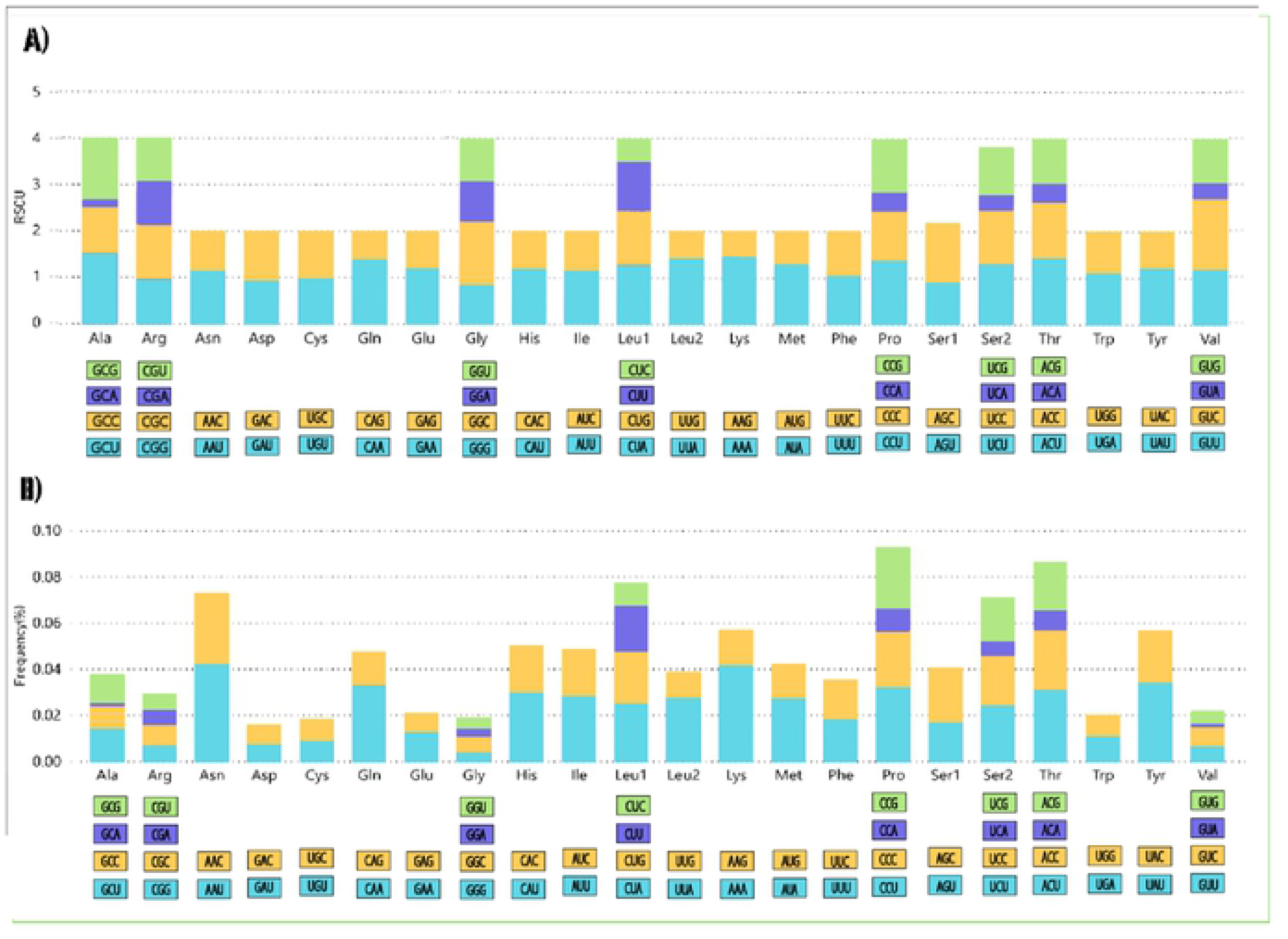
(RSCU) Relative synonymous codon usage (A) and codon usage frequency (B) ofthe mitochondrial protein coding genes of the Bangladeshi *Bos frontalis*. The stop codon is excluded. On both the figures different colors on the X-axis represent the triplet codon families corresponding to the below amino acids whereas the RSCU and codon usage frequency are plotted on the Y axis offigure (A) and (B) respectively.

### 3. 3. Transfer RNA (tRNAs) and Ribosomal RNA (rRNAs)

In the *B.frontalis* mitogenome, total of 2692bp long 22 tRNA genes are present, each of which range in size from 60 bp (trnS1) to 75 bp (trnL2) which. Most of the tRNA genes (14) are encoded by the H-strand (trnF, trnV, trnL2, trnI, trnM, trnW, trnD, trnK, trnG, trnR, trnH, trnS1, trnL1, trnT) while the remaining 8 tRNAs (trnQ, trnA, trnN, trnC, trnY, trnS2, trnE and trnP) are encoded by the L-strand (Table. 1). All tRNA sequences and structures are determined using tRNAscan-SE and MITOS [5,7]. All the tRNA genes display the typical cloverleaf secondary structure except trnS1 and trnK lacking a stable dihydrouridine arm loop (Fig. 6). Generally, such abnormal tRNA structures have been noticed in mammals including previously reported Indian gaur and Mithun [11,12,18]. Besides the standard Watson-Crick base pairs, a total of 9 unmatched base pairs are detected in these tRNAs (Fig. 6). 5 of them are G-A (two) and C-A(three) pairs, known as non-canonical pairs forming weak bonds in tRNA secondary structures. The remaining 4 mismatches include one C-U and three U-U pairs. The AT and GC skews are positive for all the tRNAs, which reveal a higher compositional count of adenine and guanine than their respective complementary nucleotides (Fig. 4). There are two rRNA genes, a 1570bp 16S rRNA gene (rrnL) and a 956bp 12S rRNA (rrnS) gene which summed up to a total length of 2526 bp. Both rRNA genes are located on the H (+) strand. The 16S rRNA subunit is located between tRNA-L2 and tRNA-V, while the 12S rRNA gene is between tRNA-V and tRNA-F (Table. 1; Fig. 2). The nucleotide composition of both rRNAs exhibits bias-ness towards AT content as well as shows positive AT skew and negative GC skew, which implies a relatively higher occurrence of adenine and cytosine than thymine and guanine in the rRNAs (Fig. 4).

**Figure 6.**
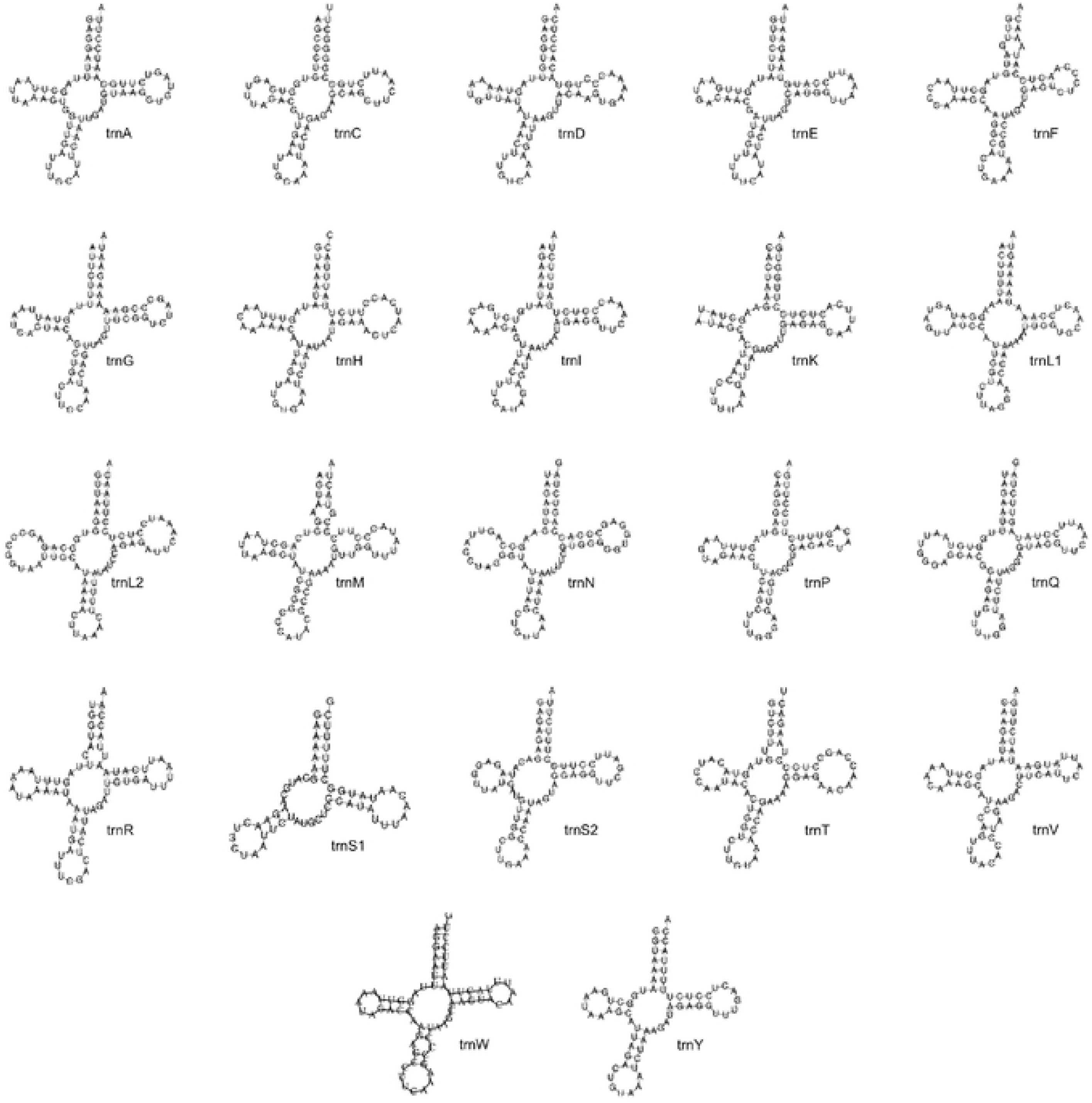
Predicted secondary structures of the 22 tRNA genes in Bangladeshi *Bos frontalis.* All tRNA sequences and structures are determined using tRNAscan-SE and MITOS [7],[5].

### 3. 4. Phylogenetic relationship

The phylogenetic analysis was performed by taking 26 bovine species belonging to 8 congeneric species like *B. frontalis, B. gaurus, B. javanicus, B. indicus, B. primigenius, B. taurus, B. grunniens, B. mutus*. In this study, our sequenced species *B. frontalis* clustered with Mithun and Gaur. This congregation of Bangladesi *B. frontalis* with Indian mithun (MK279401.1), Chinese mithun (MF614103.1) and Cambodian gaur species signifies a very close genetic relationship between mithun and gaur (Fig. 7). This finding strongly supports the unequivocal concept of the mithun being a direct descendent of gaur, which is strengthen by the results of previously reported several studies using Cytochrome b gene [19–20], 16S rRNA gene [21], SNP genotyping [22],whole mtDNA [3] and Y chromosomal DNA [23] markers. On the other hand, one of the Mithuns has paired with the *B.javanicus* (AB915322.1), a Southeast Asian cattle species commonly known as Banteng or Tembadau. The dispersed grouping of Banteng with *B.frontalis, B.gaurus* and *B.taurus* respectively indicates it’s hybrid nature likewise reported earlier [24]. On the contrary, another Chinese *Bos frontalis* (MF959941.1) has grouped with *B.indicus, B. primigenius* and *B. taurus* respectively, which exhibits the close genetic relationship between these species. Similar findings have also been obtained in previous studies using different markers [3, 25]. These outcomes demonstrate the occurrence of hybridization between domestic mithun and cattle [21,25–26] and this phenomenon is common in China. The phylogenetic relation was inferred by using the Maximum Likelihood method and Tamura-Nei model [27]. The evolutionary analyses are conducted in MEGA X [9] while the bootstrap consensus tree was inferred from 1,000 replicates is taken to represent the evolutionary history of the taxa analyzed. In this iteration process, the associated taxa that is clustered together in the bootstrap test is shown in the same clade.

**Figure 7.**
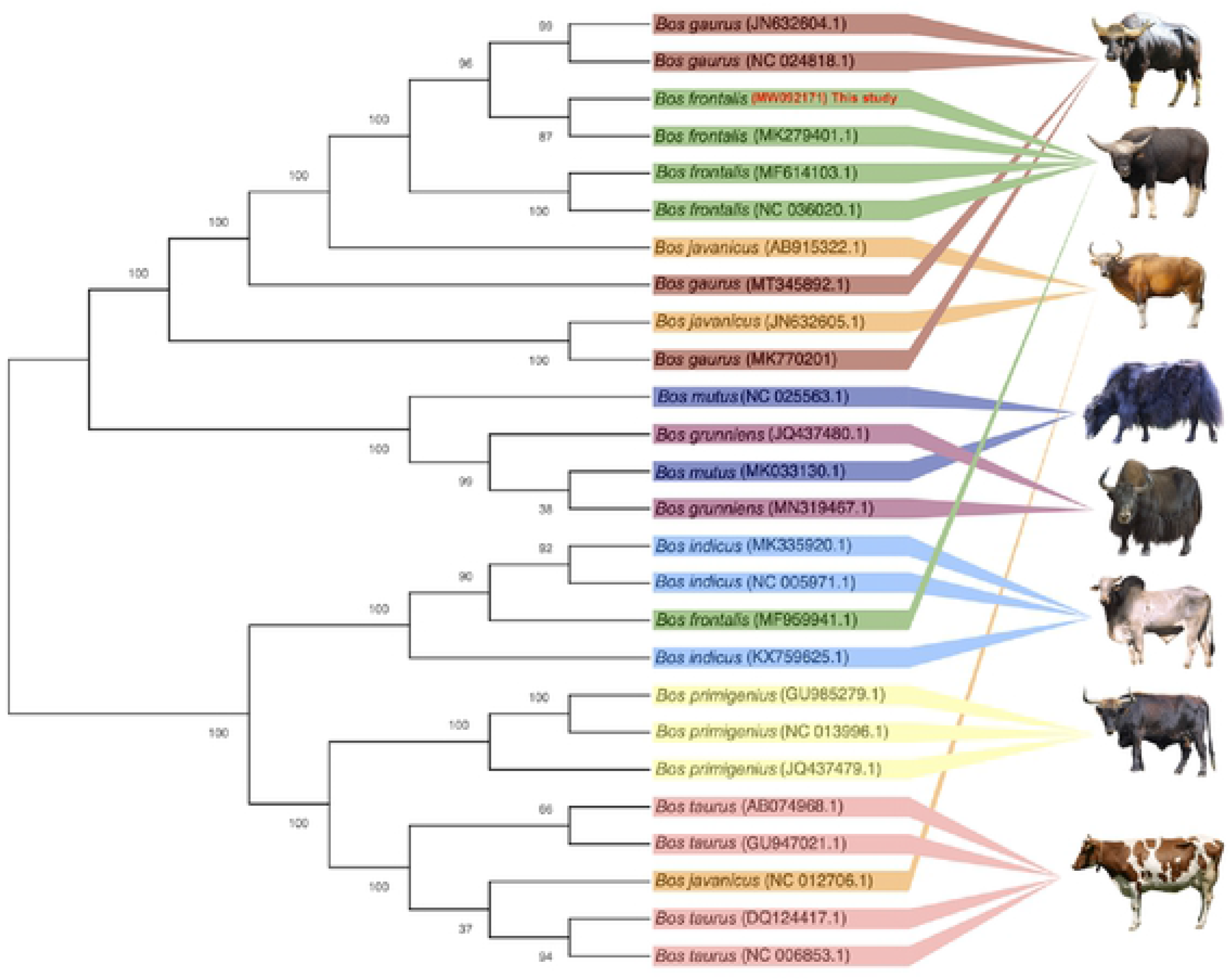
Phylogenetic relationship of 8eight *Bos* species based on their complete mitochondrial genome. Mitochondrial genome sequences are retrieved from the NCBI database except for the Bangladeshi *Bos frontalis*. The tree is generated using maximum likelihood method in MEGA X [9] and tree structure is validated by running the analysis on 1000 bootstrap iterations The numbers above the branches indicate the bootstrap support values.

## 4. Conclusion

To conclude, this study provides complete information about the mitogenome of a *Bos frontalis* that has been collected from the Bandaran Hill tract of Bangladesh. The findings of this study will pave the way for future research by animal geneticists and evolutionary biologists. This data generated will serve as a multipurpose genetic resource for further taxonomic, evolutionary, and phylogenetic analyses among closely related species. Furthermore, as closely related to Chinese gayal, Indian gayal, and Mithun, this unique resource of Bangladesh demands pertinent conservation strategies and effective implementation.

## Additional information

**Funding:** We are grateful to Bangladesh Livestock Research Institute (BLRI) to support this study financially.

